# Heritable genetic variants in key cancer genes link cancer risk with anthropometric traits

**DOI:** 10.1101/827634

**Authors:** Matteo Di Giovannantonio, Benjamin H.L Harris, Ping Zhang, Isaac Kitchen-Smith, Lingyun Xiong, Natasha Sahgal, Giovanni Stracquadanio, Marsha Wallace, Sarah Blagden, Simon Lord, David A. Harris, Adrian L. Harris, Francesca M. Buffa, Gareth Bond

## Abstract

Inherited genetic variants in tumour suppressors and oncogenes can increase the cancer risk, but little is known about their influence on anthropometric traits. Through the integration of inherited and somatic cancer genetic data, we define functional single nucleotide polymorphisms (SNPs) associated with cancer risk and explore potential pleiotropic associations with anthropometic traits in a cohort of 500,000 individuals. We identify three regulatory SNPs for three important cancer genes that associate with both anthropometric traits and cancer risk. We describe a novel association of a SNP in TP53 (rs78378222) with height, lean body mass measures and basal metabolic rate, as well as validating its known associations with brain and non-melanomatous skin cancer susceptibility. Our results clearly demonstrate that heritable variants in key cancer genes can associate with both differential cancer risk and anthropometric traits in the general population, thereby lending support for a role of genetics in linking these human phenotypes.

## Introduction

Height and other anthropometric measures have been consistently found to associate with differential cancer risk^1, 2, 3^. However, both genetic and mechanistic insights into these epidemiological associations are notably lacking^2, 4^. Increased height has been seen to associate with a number of malignancies including skin, breast colon, rectum, endometrium, ovary, kidney cancers as well as Hodgkin’s lymphoma and leukaemia^2, 5, 6^. Current theories underpinning links between increased height and increased cancer risk centre upon the role of growth hormone, insulin-like growth factor and/or insulin being pro-tumorigenic by up-regulating the Ras-MAPK and PI3K pathways leading to increased cellular proliferation^7, 8^. It has also been proposed that increased height might correspond to increased cellular number and therefore increased probability of malignancy, simply by chance^9^; however, such allometric scaling of body mass only explains part of the observed effect and it is not observed among mammal species of different size (Peto’s paradox)^10^. Like height, increased body mass index (BMI), a surrogate measure of obesity based on height and weight, has been associated with an increased risk of number of cancers including post-menopausal breast cancer, colorectal cancer and renal cancer^3, 11, 12^. Increased BMI within individuals has been mainly attributed to increasing accumulation of fat mass. It is hypothesised that adiposity causes a state of systemic inflammation, a shift of metabolite and adipokine release, and an increase of circulating insulin, secondary to insulin resistance^13, 14^. This combination may increase cancer risk. However, the exact and predominant mechanisms of how high BMI is linked with increased cancer risk are not understood. Thus, links between height, anthropometric measures and cancer remain an intense area of research.

Anthropometric traits are largely determined by genes that control cellular proliferation, metabolism and apoptosis: these attributes are also required for immortalisation of cancer cells and development of tumours ^15, 16^. It is well established that germline mutations within tumour suppressor genes and oncogenes, affect cancer risk^17, 18^; however, their influence on anthropometric traits is not well-known. From studying mouse models and rare diseases, there are indications that mutations in such genes may influence body mass composition^19, 20, 21, 22, 23^. For example, the tumour suppressor gene ARH1 inhibits cell growth. A deletion in ARH1 is commonly associated with breast and ovarian carcinoma while mice that are engineered to overexpress ARH1 are significantly smaller than wild type counterparts^24^. Mice possessing knockouts of genes within the tumorigenic hypoxia inducible pathway vary in size and respond to high-fat diets differently from those without the mutations^25, 26^. TP53 is the most frequently mutated gene in human cancers and is a key regulator of a number of cellular activities which prevent tumorigenesis, including maintaining genomic stability, controlling cell growth and metabolism^19, 27, 28^. Mouse models of TP53 mutations have demonstrated that reduction of p53 activity can increase cancer risk, alter metabolism and influence obesity in a complex and signal dependent manner^20, 21, 22, 23^.

Further evidence for the relation between body size, genetics and cancer comes human genetic disease, such as Turner syndrome. Turner Syndrome is defined by complete or partial chromosome X monosomy and is associated with a distinct clinical phenotype, including gonadal dysgenesis, cubitus valgus and short stature^29^. Short stature within this condition has been attributed to haplo-insufficiency of the SHOX tumour suppressor gene^30^. Deletions and mutations in SHOX have also been identified as an explanation for short stature within the general population^31^. Another example comes from patients with Simson–Golabi–Behmel syndrome (SGB), caused by mutations in the GPC3 tumour suppressor gene. Patients with SGB tend to have pre- and post-natal overgrowth and thus ultimately taller stature as well as an increased risk of embryonic tumours. In line with this, GPC3-null mice also display overgrowth^32^. Effects in both humans and mice are thought to be due to unshackling of the hedgehog pathways, and increasing cellular proliferation^33, 34, 35^.

These rarer diseases provide evidence that major differences in the expression of cancer genes can influence both anthropometric traits and cancer risk in specific cohorts of patients. However, it is less clear if such associations occur in the broader population. The availability of large phenotypic and genetically linked datasets, has allowed unsupervised approaches, such as genome-wide association studies (GWAS) of common single nucleotide polymorphisms (SNPs), to be deployed to further understand the link between genetics and anthropometric traits. Pleiotropy is the phenomenon whereby a single SNP or genetic mutation can influence multiple traits. Until now, it has been difficult to look for pleiotropy between functional SNPs in genes associated with cancer risk and anthropometric traits, as very few cohorts possess both comprehensive genetic data and detailed anthropometric data in the same population. In fact, the UK Biobank provides a unique opportunity to investigate such pleotropic associations in a large prospective cohort of over 500,000 participants.

## Results

Through integration and curation of the GWAS catalog, eQTL databases and the Cancer Gene Census we identified 100 SNPs, which have been shown to associate with a differential risk of developing a total of 21 different cancer types and differential gene expression in at least one tissue type of *(i)* proto-oncogenes (8 genes) *(ii)* oncogenic fusion proteins (8), *(iii)* tumour suppressors (16) and *(iv)* 15 genes that span two or more of these groups (supplementary tables 1 and 2). We define these SNPs as *cancer eSNPs*. In our analyses of the UK Biobank cohort, we found 13 cancer eSNPs associated with differences in anthropometric traits and 31 with differential cancer risk. Interestingly, 7 of these SNPs overlapped and thus displayed some level of pleiotropy between cancer risk and anthropometric traits (Fig. 1, supplementary tables 3&4). These 7 cancer eSNPs are found on three different chromosomes and in linkage disequilibrium (see methods) with at least one other identified cancer eSNP (Fig. 2). They have been found to be associated with differential expression levels of *(i)* Fanconi Anemia, complementation group A, FANCA [rs1805007, rs258322], *(ii)* tumor suppressor p53, TP53 [rs78378222, rs35850753, rs8753], and Mitogen-Activated Protein Kinase 3, Kinase 1 [MAP3K1: rs889312, rs1862626]). Both cancer eSNPs associated with FANCA expression reside in neighbouring genes; rs258322 is a non-coding variant within CDK10 and rs1805007 is a missense variant within MC1R. eSNPs associated with MAP3K1 expression are found in an intergenic region close to ZNF296 and MAP3K1 (5q11.2). In contrast, the two cancer eSNPs associated with differential TP53 expression are found in untranslated regions of the TP53 gene itself. rs78378222 is found in the 3’-UTR of TP53 and rs35850753 in is found the 5’-UTR of the d133 isoform of TP53. The third TP53 eSNP is in the 3’-UTR of the neighbouring POLR2A gene. The cancer eSNPs that showed the strongest association with anthropometric measures and cancer risk within their loci (lead cancer eSNP) were rs1805007 (FANCA, C>T, minor allele frequency in UK Biobank ^36^= 0.102), rs78378222 (TP53, T>G, MAF= 0.012) and rs889312 (MAP3K1, C>A, MAF= 0.284).

**Fig. 1:**
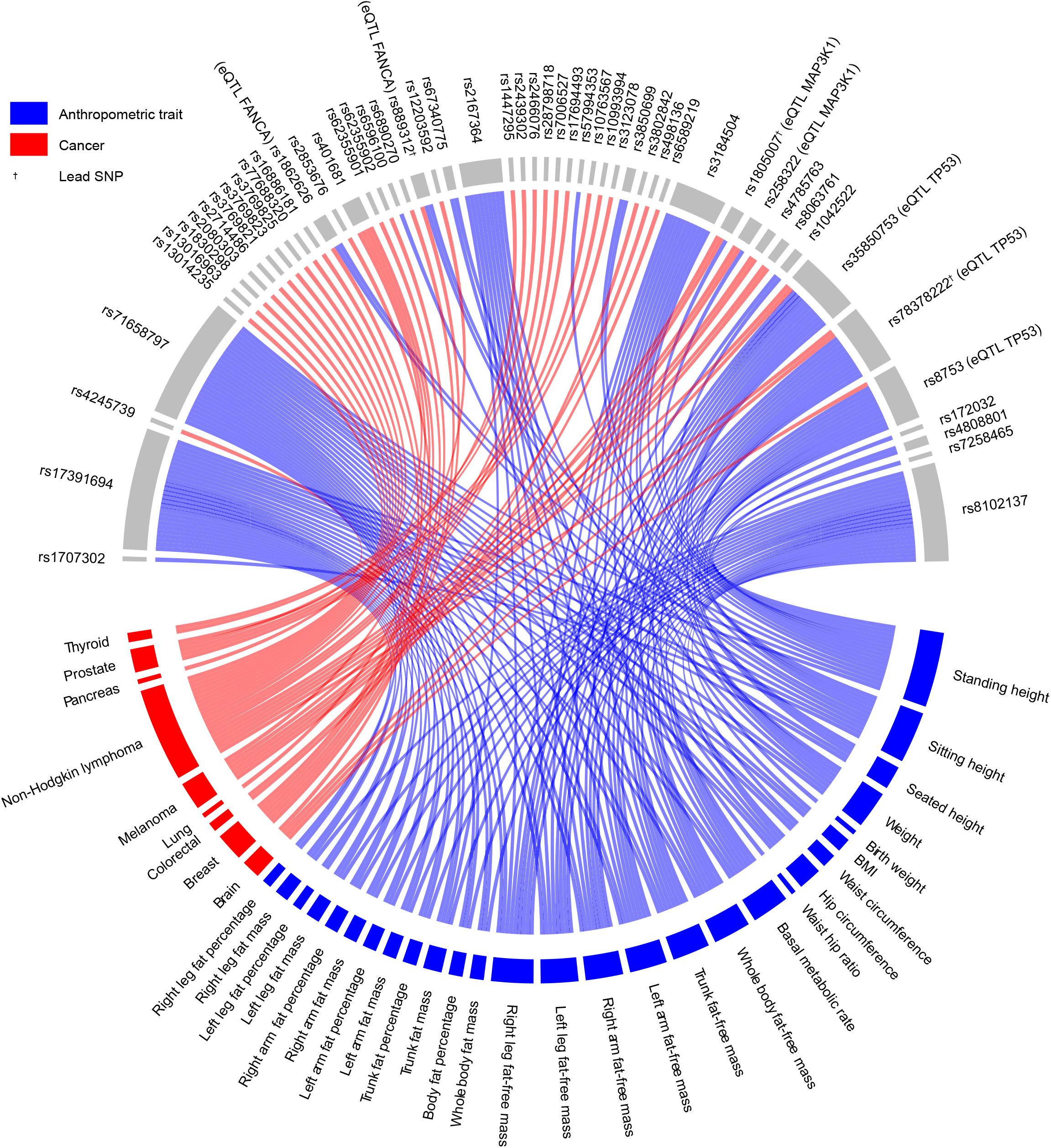
A circos plot indicating the functional SNPs that significantly associate with anthropometric traits and cancer risk. SNPs are located on the top half of the plot whilst cancer types and anthropometric traits are on the bottom half. Significant associations (Bonferroni corrected p-values <1E^−5^) are shown as solid lines. Blue lines indicate significant associations with anthropometric traits. Red lines depict significant associations with differential cancer risk.

**Fig. 2:**
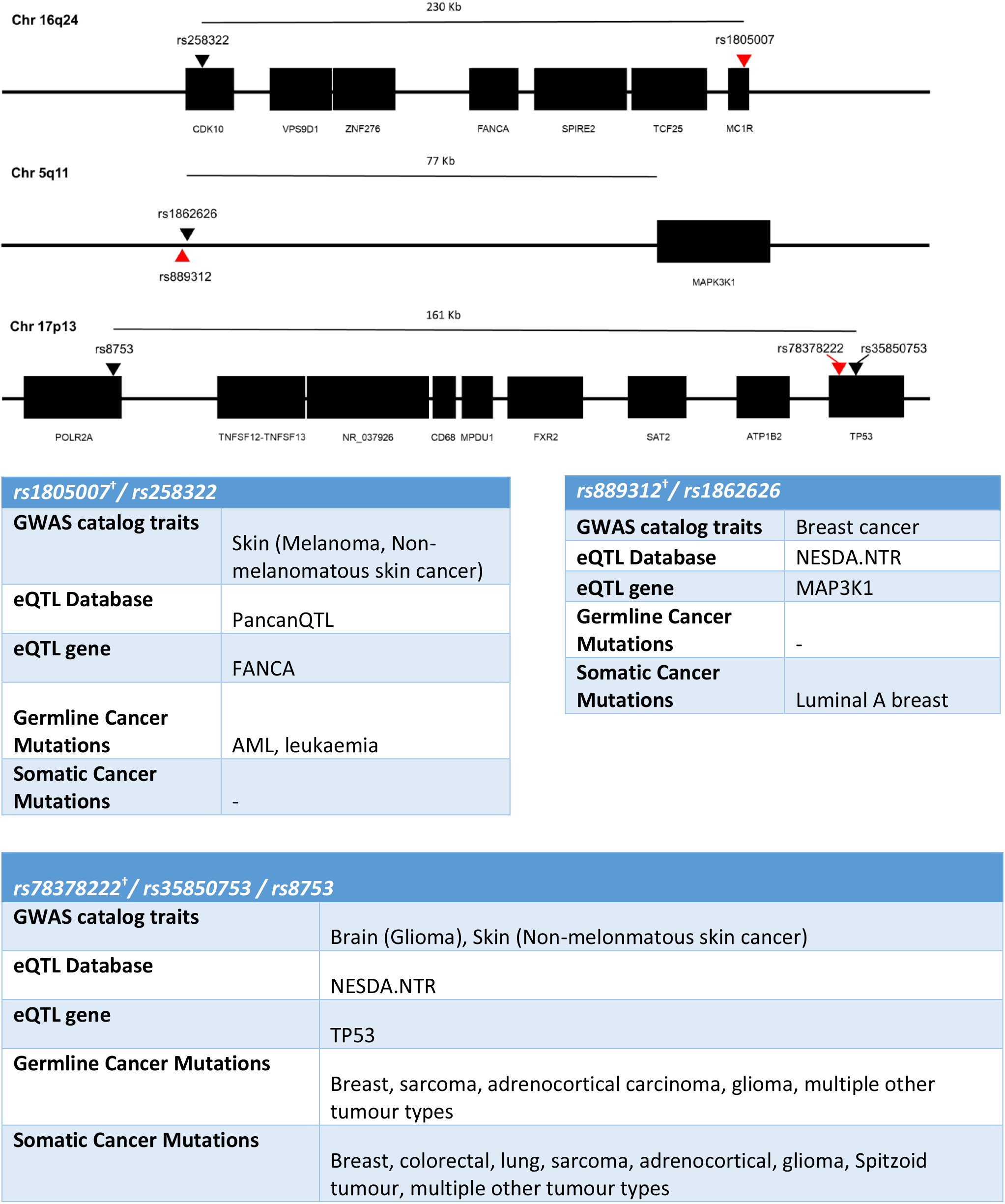
A gene map of the chromosomal regions containing the pleiotropic functional SNPs. Lead SNPs are highlighted in red. Tables summarise salient GWAS catalog, eQTL, and Caner Gene Census information for each variant.

In the UK Biobank cohort, the minor allele of rs1805007 (FANCA) was associated with an increased risk of melanoma (OR=1.63 [1.52-1.75], adjusted-p=2.57E^−41^) and non-melanomatous skin cancer (OR=1.36 [1.31-1.41], adjusted-p=7.62E^−63^). The minor allele of rs78378222 (TP53) was associated with an increased risk of brain malignancy (OR= 3.12 [2.22-4.37], adjusted p= 1.43E^−12^) and non-melanomatous skin cancer (OR= 1.46 [1.34-1.60], adjusted p= 5.20E^−18^). The minor-allele of rs889312 (MAP3K1) was associated with an increased risk of breast cancer (OR= 1.1 [1.07-1.13], adjusted p= 2.82E^−11^). Of note, as we selected these SNPs due to their noted association with differential cancer susceptibility in GWAS studies, our results provide an independent validation of these associations (Fig. 3a, supplementary table 5).

**Fig. 3:**
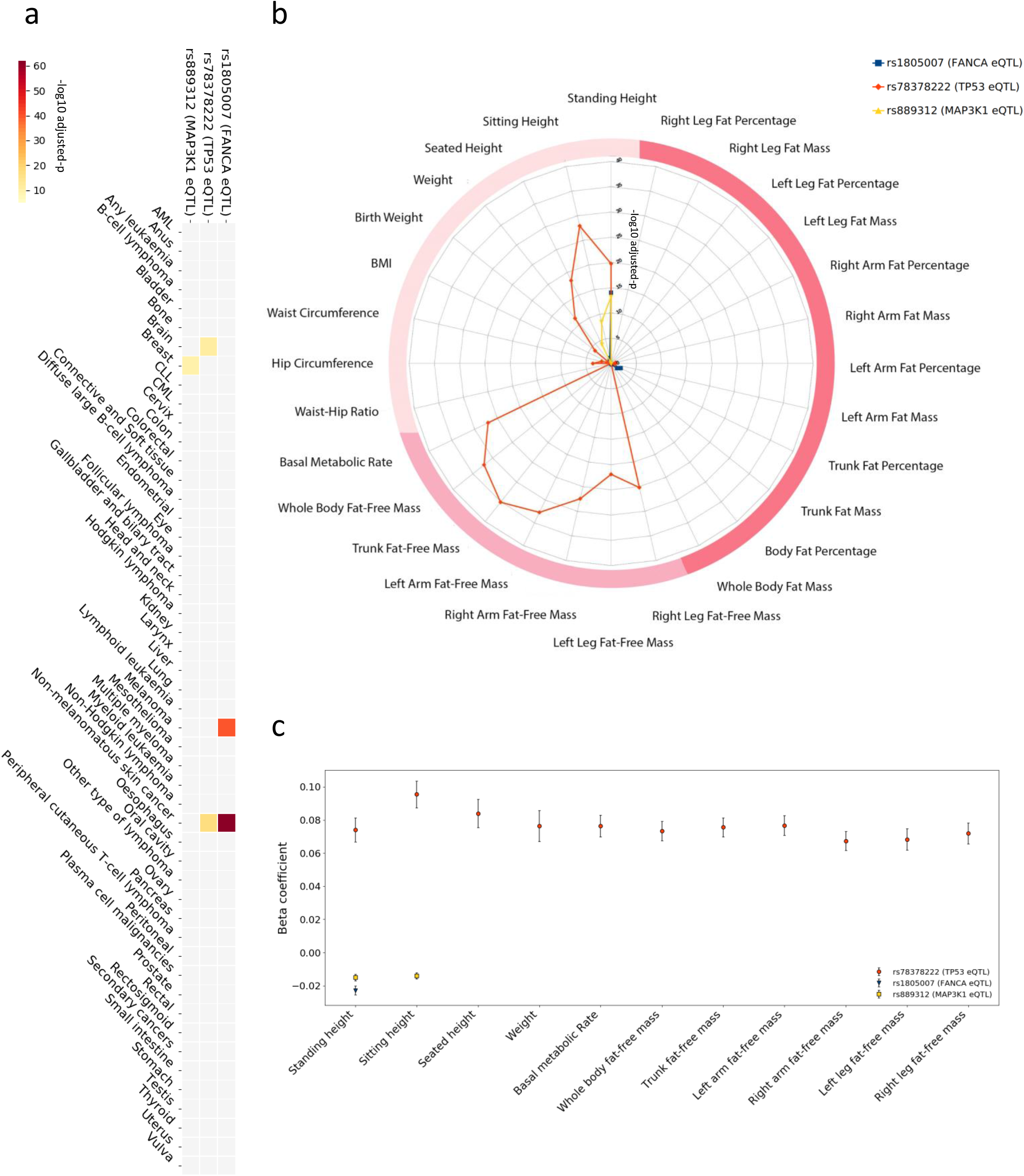
Detailed association results for cancer eSNPs with anthropometric traits and cancer risk in the UK Biobank cohort. A heatmap (a) depicting the significance of the associations of the three lead pleotropic SNPs with differential cancer risk. The colour scale represents the - log10 adjusted-p: with the darkest red on the scale being 7.62E^−63^ and the lightest yellow being 1E^−5^. A radar plot (b) illustrating the level of significance of the associations of the three lead pleotropic SNPs with anthropometric traits (radial axis: −log10 adjusted p). The darkest two pink categories are bio-electrical impedance measures (darkest being measures of fat). Traditional bedside anthropometric measures are in blush. SNPs related to FANCA (rs1805007^†^) and MAP3K1 (rs889312^†^) associate with standing height. The lead functional SNP related to TP53, rs78378222^†^, markedly associates with multiple measures of height and lean body mass, as well as basal metabolic rate. An error bar plot (c) of the beta coefficients (y-axis) of the significant associations with anthropometric traits (x-axis). Error bars denote the 95% confidence interval of the beta value.

As regards the anthropometric measures, we identified an unexpected high level of pleiotropy and very strong association between the TP53 cancer eSNPs and three measures of height and six lean body mass, as well as basal metabolic rate with p-values as low as 7.54E^−36^ (Fig. 3b). After quality control (QC) procedures, we identified 59 participants homozygous for the minor allele (increased cancer risk) of the lead TP53 eSNP (rs78378222), 9,253 heterozygous for the minor allele and 370,046 homozygous for the major allele. The minor allele carriers were on average taller, leaner and have a higher basal metabolic rate. The associations of these anthropometric traits with the minor allele of the rs78378222 SNP were markedly more significant (up to almost 3 fold) and with greater effect sizes (up to almost 5 fold) than for the other pleiotropic eSNPs (Fig 3c, supplementary table 3).

The cancer eSNPs for TP53 demonstrated strikingly strong associations with 10 different anthropometric traits in the UK Biobank cohort. In contrast, the cancer eSNPs related to FANCA [rs1805007] and MAP3K1 [rs889312] were associated with standing height but not lean body mass measures. After QC, we identified 4271 participants homozygous for the minor allele of the lead FANCA eSNP, 68,956 heterozygous for the minor allele and 306,131 homozygous for the major allele. The minor allele for the lead MAP3K1 eSNP was found in 185,184 participants (30,394 homozygous, 154,790 heterozygous). The FANCA eSNP [rs1805007] only associated with standing height (beta= −0.02 ± 0.002, adjusted-p=9.20E^−15^), while the MAP3K1 eSNP [rs889312] associated with broader range of height measures (standing height beta= −0.02 ± 0.004, adjusted p= 6.11E^−14^ and sitting height beta= −0.01 ± 0.001, adjusted p= 3.34E^−9^). We sought validation for these associations with standing height in the independent GIANT consortium dataset. To do so, we examined at all lead and linked SNPs for each pleiotropic eSNP in the results of the meta-analysis conducted by Wood et al. (2014)^37^. We were able to identify eSNPs for MAP3K1 (rs889312, rs1862626) and FANCA (rs1805007, rs258322), but none for TP53, due to the low minor allele frequency. Three of these eSNPs significantly associated with height in the Wood et al. meta-analysis: (i) rs258322 FANCA eQTL (p=1.5E−09, Beta=−0.029), (ii) rs889312 MAP3K1 eQTL (p=2.3E−08, Beta=−0.018) and (iii) rs1862626 MAP3K1 eQTL (p=3.4E−08, Beta=−0.018). Interestingly, both linked MAP3K1 eSNPs and a FANCA eSNP passed the significance threshold and, reassuringly, the directions of the allelic associations are consistent with our findings. The other FANCA eSNP only just fell short of the GWAS significance threshold (rs1805007, 4.3E−04, Beta = −0.024). Together, these data clearly link eSNPs for MAP3K1 and FANCA with height.

## Discussion

This is the first comprehensive study providing evidence that functional common genetic variants in oncogenes and tumour suppressor genes can associate with both anthropometric traits and cancer risk in the general European population and in the same cohort. SNPs exhibiting these pleiotropic associations in our study are found in three different loci: *(i)* MAP3K1 (two SNPs in linkage disequilibrium), *(ii)* FANCA (two SNPs in linkage disequilibrium) and *(iii)* TP53 (three SNPs in linkage disequilibrium). Observations gained from mouse models designed to alter signalling pathways involving *FANCA* and *TP53*, suggested that such associations with anthropometric traits in humans could be possible. For example, targeted disruption of exons of the Fanconi Anemia group A (FANCA) gene in mice results in altered anthropometric traits, including growth retardation as well as differential cancer risk^38^, ^39^. Furthermore, mouse models of TP53 mutations have clearly demonstrated that reduction of p53 activity can result in increased cancer risk, altered metabolism and influence obesity in a complex and signal dependent manner^20, 21, 22, 23^.

The strongest associations we observed with both anthropometric traits and cancer risk are loci related to TP53. p53 is key regulator of a number of cellular activities which prevent tumorigenesis including maintaining genomic stability, controlling cell growth and metabolism^27, 28^. TP53 is the most frequently mutated gene in human cancers^19^. Moreover, in all families with similar TP53 mutations in their heritable genomes, a dramatic increase in cancer risk is observed (Li-Fraumeni Syndrome, LFS)^17, 40^. In recent studies, it has been shown that LFS patients not only have an increased risk of developing cancer but also an increased capacity for oxidative phosphorylation^41^, providing a potential link with anthropometric traits and basal metabolic rate. Furthermore, the well-tolerated anti-diabetic drug metformin, which is thought to inhibit mitochondrial complex 1, increases cancer-free survival in a mouse model of LFS and reduces proliferation in cancer cell lines^42, 43^. Metformin is now being trialled in LFS patients to hopefully provide a preventative option for these high-risk patients (ClinicalTrials.gov number: NCT01981525). Based on the potential link with oxidative phosphorylation, this intervention might also be trialled in those carrying the minor allele of the TP53 cancer eSNP (rs78378222).

The TP53 mutations found in LFS are rare in the general population^44^. However, here we show relatively frequent SNPs related to p53 affect cancer risk and anthropometric traits. Notably, the allele of a SNP in the polyadenylation signal of p53 (rs78378222[C]) which is found in approximately 1% of populations of European descent, has been shown to impair 3’- end processing of p53 mRNA, resulting in a reduction of p53 protein and an increased risk for glioma and basal cell carcinoma as well as affecting head circumference and intracranial volume ^45, 46^. Here we not only validate these cancer associations in a separate cohort (non-melanomatous skin cancer, OR= 1.46 [1.34-1.60], adjusted p= 5.20E^−18^, brain malignancy, OR= 3.12 [2.23-4.37], adjusted p= 1.43E^−12^) but also show that carriers of this allele tend to be taller, leaner and have a higher basal metabolic rate (standing height, adjusted-p= 2.18E^−24^, beta= 0.073 ± 0.007, whole body fat free mass, adjusted-p= 8.34E^−37^, beta= 0.073 ± 0.005, basal metabolic rate, adjusted-p= 1.13E^−31^, beta= 0.076 ± 0.006).

Prior to this study, the association between increased height and non-melanomatous skin cancer/brain malignancy were well established, however, mechanistic explanations were lacking^2, 47, 48^. Our study proves the concept that functional loci in well-characterised tumour suppressors and oncogenes alter both cancer risk and anthropometric traits and, of particular interest, identifies a strong, new, association between the rs78378222[C] SNP in the polyadenylation site of p53 with both increased risk for developing non-melanomatous skin cancer/brain malignancy and increased height, lean body mass and basal metabolic rate, thereby offering a novel genetic link between these anthropometric traits and cancer risk.

## Methods

### The UK Biobank

The UK Biobank is a cohort of ∼500,000 UK residents who volunteered to have their clinical, lifestyle, anthropometric and genetic data collected for research. Data was collected in a number of ways, including face-to-face interview, touchscreen assessment and from centralised clinical registers (e.g. the Cancer register and Death register). Participants were aged between 40-69 years at recruitment^49^. UK Biobank obtained informed consent from all participants and all protocols were approved by the National Research Ethics Service Committee. All baseline data used in this study was collected in 22 UK centres, between 2006-2010. This study was conducted under the UK Biobank approved application (#24456, PI Gareth Bond).

### Anthropometric data

During the baseline assessment, participants had various anthropometric traits measured directly or by bioelectrical impedance. Bioimpedance data was measured using a Tanita BC418MA body composition analyser. Participants stood barefoot on the analyser and held the metal handles. This device produced measurements of fat mass, fat-free mass and basal metabolic rate. Further information on anthropometric data collected is found here^50^.

### Cancer data

Cancer occurrences were defined by presence of a cancer international classification of diseases (ICD) code in the UK Cancer register or the UK Death register. To maximise the number of individual cancers cases, we combined ICD9 and ICD10 codes of identical cancers. This was done by a clinician to ensure the matching was accurate, for instance malignant neoplasm of brain was defined as C71 (ICD10) and 191 (ICD9). With the goal of increasing power, when appropriate we merged ICD codes into clinically relevant groups. We ran individual cancers (combined ICD9 and ICD10 codes) and clinically relevant groups in our analysis.

### Genetic data

Blood samples were collected when participants were recruited, and DNA extracted^51^. DNA was then genotyped on either the Affymetrix UK BiLEVE Axiom array or the Affymetrix UK Biobank Axiom array (Santa Clara, CA, USA). Imputation was based upon a merged reference panel of ∼90 million biallelic variants, from the 1000 Genomes Phase 3^52^ and the UK10K^53^ haplotype panels. Imputation was performed using IMPUTE2 as described^54^, producing 488,295 genotyped participants.

### Sample quality control

In addition to the standard quality control, we carried out further quality control steps to ensure robustness of our analyses. We excluded individuals based on: a mismatched value between self-reported and genetic sex (data-field: 22001 and 31), level of genotype missingness of >0.05 (data-field: 22005), genetic relatedness factor with kinship coefficient of >0.0442, sex chromosome aneuploidy (data-field 22019), outliers for heterozygosity or missing rate (data-field: 22027). We selected the European population based on self-reported ethnicity (data-field: 21000) by excluding non-white ethnic background. This left a study population of 379,358 suitable genotyped individuals.

### Identification of functional cancer gene SNPs

We identified genes confirmed to be involved in carcinogenesis in the COSMIC Cancer Gene Census (release v88, 19th March 2019): the current reference record of genes containing cancer driver mutations. We then selected all SNPs annotated to these genes that have been significantly associated with differential cancer risk in genome-wide association studies in the GWAS catalog^55^. Significant associations with cancer risk were defined with a p-values of <5E^−08^cut-off. Finally, we required that these SNPs were also associated with differential expression of the cancer gene in at least one expression quantitative trait loci (eQTL) database. cis-eQTL databases utilised were GTEX, NESDA/NTR and PancanQTL^56, 57, 58^.

### SNP quality control

SNP exclusions were Hardy-Weinberg equilibrium with p-value less than 1E^−10^, a minor allele frequency less than 0.0001, level of missingness more than 0.05 or an imputation score less than 0.8 (as per http://www.nealelab.is/uk-biobank/).

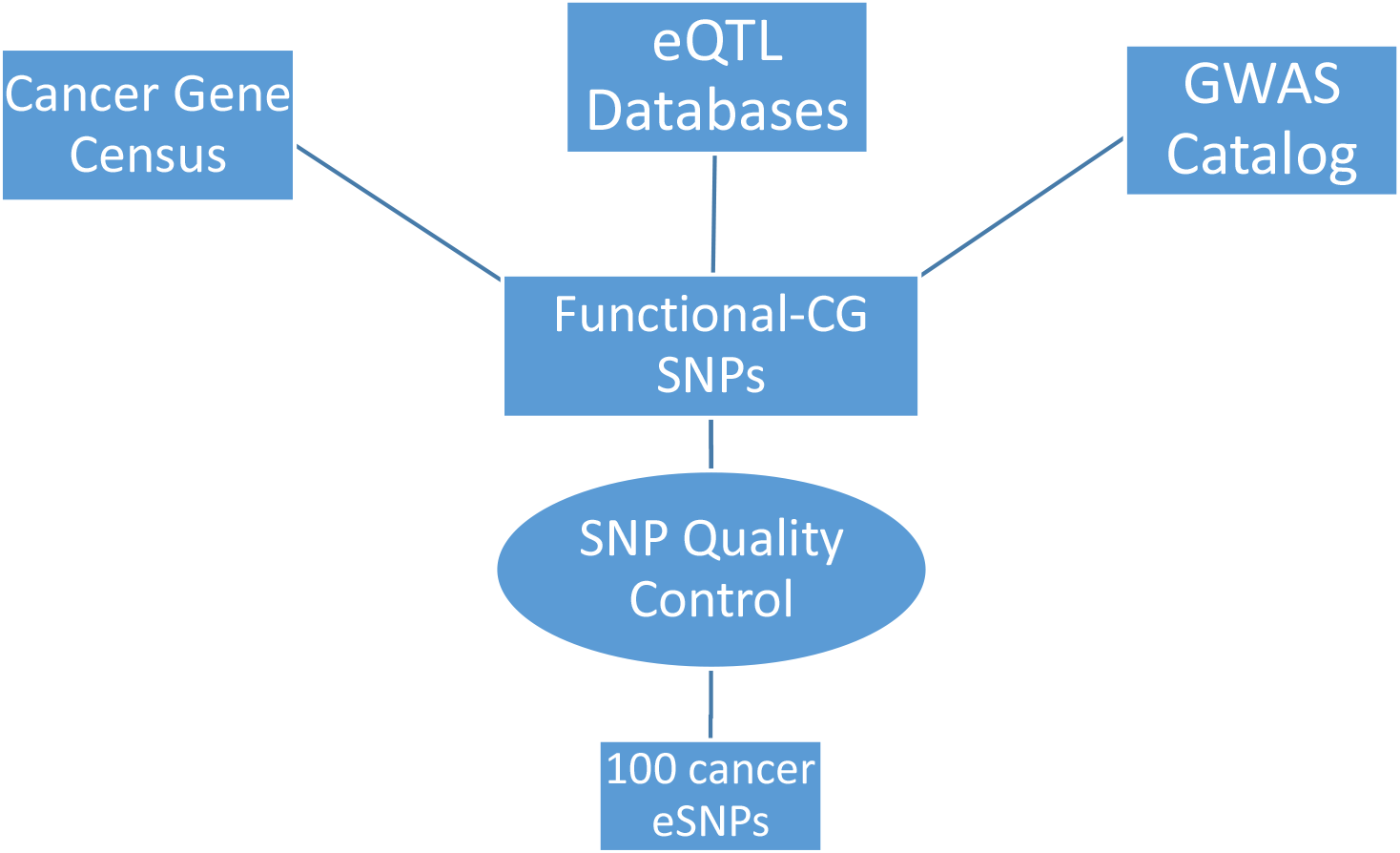

### Analysis

To carry out the SNP-wise analysis on all functional-CG SNPs we used SNPTEST (version 2.5.4) ^59^, and employed the frequentist approach under dominant, additive and recessive inheritance models, using sex, age and genetic principal components (1-20 PCs) as covariates. Genetic PCs were included as covariates to control for hidden population structure. We controlled the genotype uncertainty by implementing the missing data likelihood score test. Within SNPTEST, we used the frequentist approach under an additive inheritance model. P-values of SNP-wise association were adjusted by the stringent Bonferroni correction based on number of tested traits (28 anthropometric traits, 50 cancer types) multiplied by the number of eSNPs (100). Significant associations were defined Bonferroni correction p values below the threshold of 1E^−5^.

### Linkage and lead SNPs

R^2^ and D’ coefficients were calculated to evaluate the degree of Linkage Disequilibrium (LD) between different loci using LDlink v3.7 (D’> 0.8, R^2^>0.4 https://ldlink.nci.nih.gov/)^60^. Lead SNPs were defined as the SNPs that were most strongly associated with the traits in question for each locus^61^.

Due to the low value of r2 in some of the potential LD eSNPs we carried out a leave-one-out analysis:

– For each lead SNP:

- All participants from the cohort carrying the lead SNP were removed.
- The association for each of the potential LD eSNPs was performed (e.g. lead SNP rs783222 carriers were removed whilst rs35850753 and rs8753 were tested for their associations with cancer and anthropometric traits).
- Adjusted p-values from the whole cohort association analysis with those obtained in this analysis were compared.
- If the association of the potential LD eSNPs was not significant after the removal of lead SNP carriers (p-values>0.05), we considered the eSNP in Linkage Disequilibrium.

## Acknowledgments

Dawn O’Reilly for proofreading the article

## Footnotes

† denotes lead cancer eSNP

